# A Precuneal Causal Loop Mediates External and Internal Information Integration in the Human Brain

**DOI:** 10.1101/2020.08.20.259846

**Authors:** D. Lu, I. Pappas, D. K. Menon, E. A. Stamatakis

## Abstract

Human brains interpret external stimuli based on internal representations. One untested hypothesis is that the default-mode network (DMN) while responsible for internally oriented cognition can also encode externally oriented information. The unique neuroanatomical and functional fingerprint of the posterior part of the DMN supports a prominent role for the precuneus in this process. By utilising imaging data during two tasks from 100 participants, we found that the precuneus is functionally divided into dorsal and ventral subdivisions, each one differentially connecting to internally and externally oriented networks. The strength and direction of their connectivity is modulated by task difficulty in a manner dictated by the balance of internal versus external cognitive demands. Our study provides evidence that the medial posterior part of the DMN may drive interactions between large-scale networks, potentially allowing access to stored representations for moment to moment interpretation of an ever-changing environment.

## Introduction

The intrinsic coupling between regions in the human brain (or functional connectivity/FC) is not random but forms consistent spatial patterns known as Intrinsic (functional) Connectivity Networks (ICN) (Seeley et al., 2007). Each ICN’s relevance to cognitive function has been established from activation studies, from which we can infer what information features are encoded in different networks (Cole et al., 2014).

Interactions between ICNs are thought to be a form of information exchange serving cognitive demands, although the precise functional role of those interactions remains an active area of research (Bressler & Menon, 2010). A case in point is the interaction between the default mode network (DMN) and cognitive control networks which have been reported to be anti-correlated (Fox et al., 2005) and contraposed in their cognitive function (Weissman et al., 2006). This view however is increasingly challenged by accumulating findings. While the majority of DMN studies focus on *resting state*, i.e. in the absence of external stimulation, emerging evidence shows that the DMN is indeed engaged during goal-directed tasks (Elton & Gao, 2015; Spreng et al., 2014; D. Vatansever et al., 2015; Deniz Vatansever et al., 2015, 2018), as opposed to being a “resting-state” network.

A conciliatory model that can account for both the internal and external role of DMN is that of predictive coding. The basic assumption is that conscious processing, whether externally oriented or not, always entails some form of internal processing. Thus internal and external representations are not dissociated but rather constantly update each other in a Bayesian fashion (Barrett & Simmons, 2015; Friston, 2012). Given its multifaceted role in external and internal processing and motivated by pharmacological and brain injury studies that highlight the DMN’s prominent role in conscious processing (Liu et al., 2015; Perri et al., 2016; Vanhaudenhuyse et al., 2010), we hypothesised that the DMN is fundamental when considering the neural instantiation of such an account.

The DMN, which comprises medial frontal and medial posterior parietal cortices as well as angular gyri and hippocampus, is neither anatomically nor functionally homogeneous (Braga et al., 2013; Braga & Buckner, 2017; Kernbach et al., 2018). Among the DMN regions, the precuneus (PCu/PCC) has attracted significant interest due to its complex neuroanatomical, metabolic and functional fingerprint (de Pasquale et al., 2012; Raichle et al., 2001; Spreng et al., 2009; Utevsky et al., 2014). The PCu has been characterised as the nexus of the DMN’s integrative fingerprint (Utevsky et al., 2014). It is engaged in a broad range of cognitive tasks, including both internal representation and externally-oriented, goal-directed tasks (Cavanna & Trimble, 2006; Fletcher et al., 1995) and its activity/connectivity can discriminate between conscious/unconscious states (Utevsky et al., 2014). It has exceptionally high metabolic rate (Raichle et al., 2001) and is suggested to have undergone the biggest brain morphological expansion through human evolution (Bruner & Iriki, 2016). The PCu is also a major connectivity hub from a graph-theoretic perspective (P van den Heuvel & Sporns, 2011; Tomasi & Volkow, 2011; van den Heuvel & Sporns, 2013). Based on its functional features we hypothesised that the PCu may play a key role in integrating external information with internal representations.

To investigate this claim, we first ascertained the DMN’s involvement in cognitive tasks by employing a NeuroSynth meta-analytic framework. We then used MRI datasets (diffusion MRI, resting-state & task-state fMRI) from the Human Connectome Project to investigate activity as well as functional and structural connectivity of the PCu with whole-brain ICNs. We found that the dorsal and ventral PCu subdivisions had distinct activity and connectivity patterns modulated by task difficulty, a factor that influenced a mirrored interplay between the PCu and internally/externally oriented networks. This feature of the PCu suggests that it might support the integration between internally and externally oriented cognitive processes. Moreover, Dynamic Causal Modelling (DCM) provided evidence for directed couplings between the two PCu subdivisions, hinting at a combinatorial processing mode where incoming information is associated with internal representations.

## Results

### The DMN during tasks: A meta-analytic perspective

We first validated that the DMN is indeed involved during goal-directed tasks, as findings regarding DMN’s involvement during tasks are still disputed (Fransson, 2006). We conducted a meta-analysis with the latest NeuroSynth database, using a text-based filter (“attention* or execut*”) to search for tasks that require either attention or executive processing or both (since the two cognitive processes are hardly separable in practice), which yielded 1219 fMRI studies. Using the *forward-inference* estimation, we found that a large portion of the DMN (1667 voxels) was associated with these tasks (Figure 1a), with significant clusters (*Z* > 4) located at the PCu and angular gyrus (AnG). In contrast, the *reverse-inference* estimation only revealed 68 voxels in DMN regions (Figure 1b). We acknowledge that NeuroSynth meta-analyses do not differentiate between activation and deactivation but based on previous literature which has shown PCu and AnG involvement in cognitive processing, we believe what we see here is that the tasks employ posterior DMN regions, yet their activation is so pervasive among all kinds of tasks that cannot be exclusively associated with the specified goal-directed tasks. We propose that the meta-analysis indicates a domain-general role for posterior DMN areas which may serve cognitive demands by providing contextual information.

**Figure 1:**
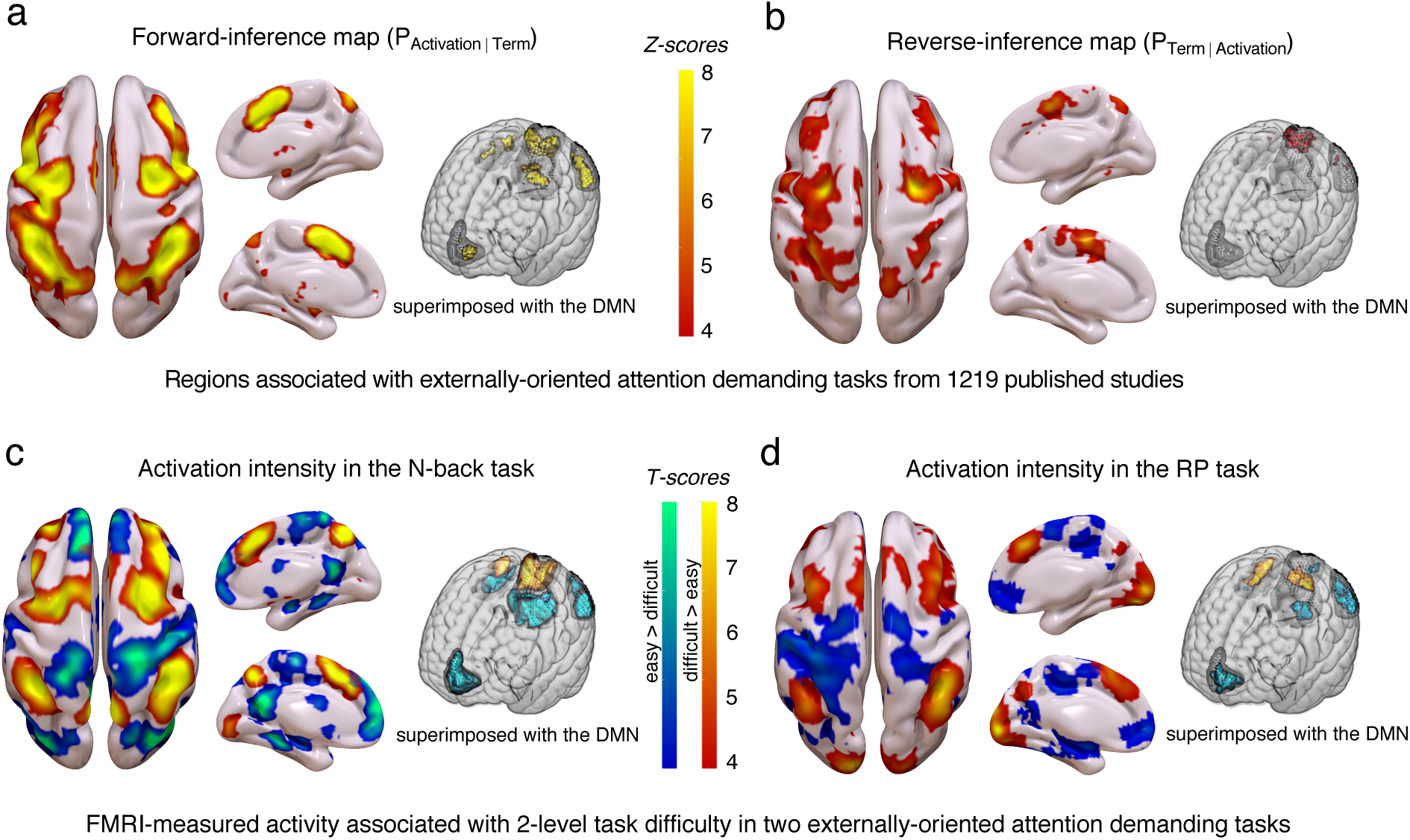
Meta-analysis and univariate analysis shows DMN engagement during the two tasks. (a) and (b) demonstrate the NeuroSynth meta-analysis results, using “attention*” or “execut*” as keywords to search for goal-directed tasks that require attention and executive function. (a) Forward inference shows that DMN subregions are active in the tasks. The forward inference map is produced by calculating the convergence of brain regions most consistently activated by certain cognitive processes. (b) Reverse inference shows that not many DMN regions are specifically associated with these tasks. The reverse inference map is calculated as the likelihood of a search term being used in a study given the presence of reported activation, and it reflects the brain activation specific to a certain cognitive process (Yarkoni et al., 2011). (c) and (d) show that activation of posterior DMN regions is associated with the N-back and RP task (from the HCP database). Warm/cold regions in the brain heatmap indicate higher/lower activity in the difficult condition compared to the easy condition. For panels (a) and (b) standardised Z scores and for panels (c) and (d) T-scores indicating activation strength are provided with colour scales. For highlighting activation in relation to the DMN, 3D renderings of the DMN are shown in shaded grey, on which our activated regions are superimposed. This was constructed by superimposing the Z-score maps (with the cut-off of 3) and T-scores (of significant clusters with P_cFWE-corr_ < 0.05), with the DMN atlas (as defined by the *Conn* network atlas).

To investigate the relationship between DMN and other ICNs in more detail during task execution, we selected two fMRI paradigms (100 young healthy participants) from the HCP dataset. The tasks we chose were the N-back working memory task and the Relational Processing (RP) task, which have attention-demanding, goal-directed features, and engage similar cognitive domains at two difficulty levels (N-back: *2-back* > *0-back* conditions, RP: *relational processing* > *matching* conditions); therefore, the two tasks afforded us the possibility of inferring the brain’s response to varying cognitive demands regardless of the specific cognitive content. The merit of using *two* tasks is to demonstrate that the DMN’s function is generalisable. As a sanity check for both tasks, the accuracy rate (N-back task: t = 2.44, p = 0.016; RP task: t = 8.80, p = 1.012e-15) was significantly higher in the easy than difficult conditions, and reaction times (RT) were significantly shorter (N-back task: t = 10.58, p < 2.2e-16; RP task: t = 10.02, p < 2.2e-16) (Table 2), suggesting that the cognitive load in difficult conditions was indeed higher than in easy conditions. To avoid task-irrelevant cognitive confounds, we only included trials with correct responses for further statistical analyses.

Since the meta-analysis results indicated a domain-general role for the DMN, we first examined the activated regions for the two HCP goal-directed tasks to confirm our hypothesis that the DMN (especially posterior) could be activated by attentional-demanding and goal-directed tasks. For the difficult versus easy contrast, most of the task-induced activations were in cognitive control regions (Figure 1c,d; activation tables are presented in the Supplementary Materials), in line with the NeuroSynth meta-analysis and previous studies using similar tasks. However, we also found significant activations in the AnG and the PCu, parts of the DMN according to the two independent brain atlases we adopted in this paper (see Methods). For the easy versus difficult contrast, wide-spread deactivations were found in the DMN. However, significant deactivation clusters were also found in non-DMN regions.

Overall, our task activation studies indicated that the PCu was differentiated into dorsal and ventral parts (dPCu, vPCu) (Table 1, Figure 2a), which were respectively more active during higher and lower cognitive demand conditions. This d/vPCu functional division was shared by the two tasks.

**Table 1:** The N-back and the RP task resulted in the same pattern of PCu fragmentation. The dorsal PCu (dPCu) and ventral PCu (vPCu) were identified as the regions responsive to different cognitive load: the dPCu was activated while the vPCu was deactivated by an increased cognitive demand in difficult versus easy conditions. The N-back task provided a statistically stronger demonstration (larger significant clusters) of this, possibly because the task had more trials i.e. bigger sample size (80 trials in the N-back, vs. 27 in the RP task).

|  | dPCu |  |  | vPCu |  |  |
| --- | --- | --- | --- | --- | --- | --- |
| | coordinate | cluster size | $P_{\text{cFWE-corr}}$ | coordinate | cluster size | $P_{\text{cFWE-corr}}$ |
| RP task | -4 -72 38 | 241 | 0.002 | -8 -60 18 | 216 | 0.005 |
| N-back task | 8 -66 52 | 2175 | 0.000 | -8 -60 14 | 1547 | 0.000 |

**Figure 2.**
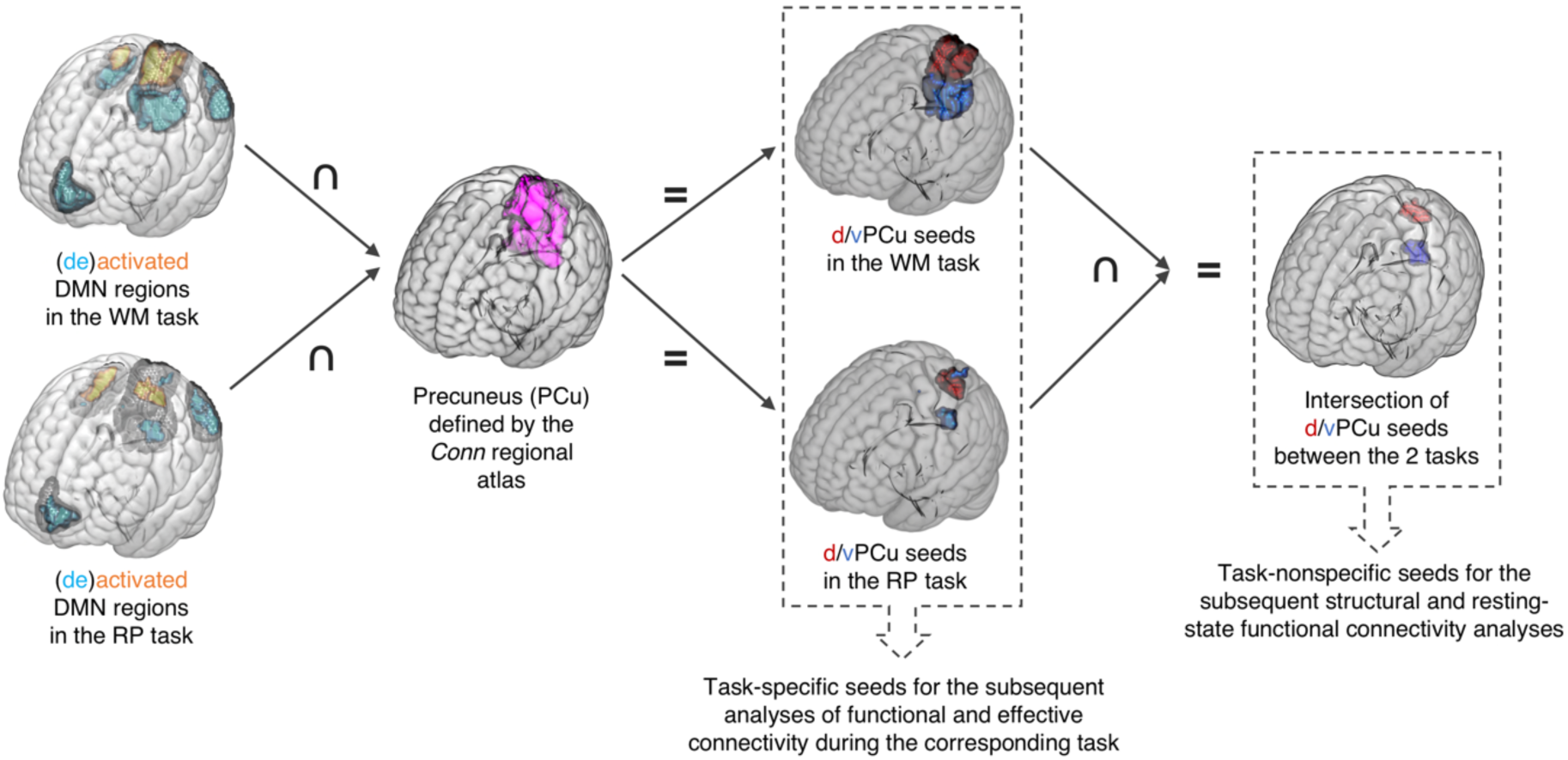
Seed derivation for the connectivity analyses. Task-specific seeds for functional and effective connectivity during tasks were derived by overlapping activation maps from either the N-back or RP task, and the DMN and PCu masks from the Conn atlas. Analysis for each task was carried out independently. For task-independent structural and resting-state functional connectivity, the PCu seeds (d/vPCu) were defined as the spatial intersection of the activated or deactivated clusters in both tasks, the DMN and the PCu masks.

### Differential connectivity between v/dPCu and ICNs

To explore further the differential response of the dorsal and ventral aspects of the PCu to task demands and the role they may serve in processing internal and external information, we next focused on the d/vPCu’s connectivity, including structural, resting-state, task-state and effective connectivity. We demonstrated that there is a disparity in the v/dPCu’s structural connectivity (SC) and resting-state (rs) FC (connectivity results for each seed are presented in Supplementary Materials) and the difference follows the pattern of internally and externally oriented cognitive function. Specifically, by comparing the vPCu and dPCu’s whole-brain SC, we found that the vPCu is more connected with the vmPFC, ACC and hippocampus, which are often implicated in value encoding, emotion, interoception and episodic memory (Chudasama et al., 2013; Euston et al., 2012; Gu et al., 2013; Pessoa, 2008), while the dPCu is more connected with regions in cognitive control networks that are associated with executive, attentional control and goal-directed behaviour (Figure 3a, 3b). We also found similar differentiation in rsFC of the d/vPCu.

**Figure 3:**
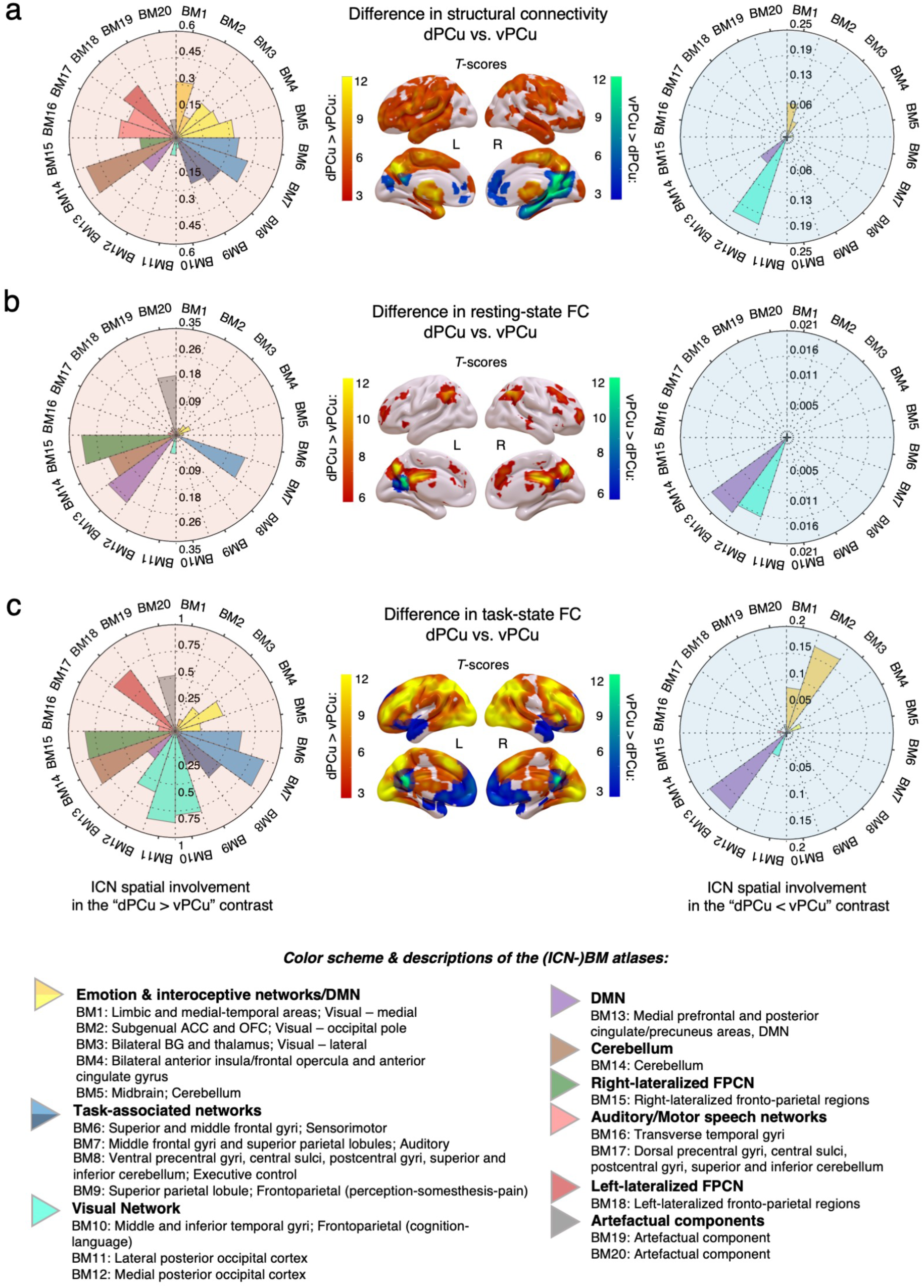
(a), (b) and (c) indicate whole-brain contrasts of the connectivity between dPCu and vPCu. Structural connectivity is in (a), resting-state FC in (b) and task-state FC in (c). Task-state FC was calculated as a partial correlation, controlling for the effect of event-related BOLD signals (i.e., disregarding the apparent correlation caused by stimulus-driven activity). T scores for the statistical effect of the connectivity difference are mapped out on the 3D brain reconstructions. Hot colours represent areas that demonstrate stronger connectivity with the dPCu than with the vPCu, and vice versa for the cold colours. Circular wedge plots to the left and right are a representation of ICN spatial involvement i.e. show voxel overlap between canonical ICNs and the connectivity results. The canonical ICNs are defined by the BrainMap/BM ICN atlases from the *ICN_atlas* toolbox (Kozák et al., 2017). Cognitive domains and descriptions of the BM/ICN atlases for the ICNs are also provided. The same colour scheme for the (ICN-) BM atlases was used throughout the paper.

To establish whether the v/dPCu connectivity differentiations in physical wiring and resting state also have functional significance, we next investigated overall FC patterns during tasks. We achieved this by computing the v/dPCu’s endogenous task-state FC (tsFC), i.e., the correlation of timeseries throughout the course of the two tasks, after regressing out the event-related haemodynamic response function (HRF). Again, contrary to the well-known DMN “anti-correlation” argument, we found a large cluster of positive tsFC with the d/vPCu, centring at the seeds themselves, and covering regions within and beyond the DMN (See supplementary materials for the composite of network domains of the v/dPCu’s tsFC). Overall, the PCu’s tsFC with a wider range of regions, including ones outside the DMN, is probably an indication of more intensive cognitive computation during tasks.

To summarise, thus far we have presented evidence from SC, rsFC and tsFC showing that the dorsal and ventral PCu are differentially connected with internally-oriented networks (IoN) and the externally-oriented networks (EoN).

### Cognitive demands modulate the effective coupling between the v/dPCu and internally/externally oriented networks

The connectivity profiles of the d/vPCu as established so far suggest that the PCu overall has the structural framework necessary and may functionally serve as a platform connecting internal and externally related information. To confirm that this is cognitively relevant we investigated whether d/vPCu connectivity during tasks is modulated by cognitive load and whether this modulatory effect can predict task performance. We therefore investigated the Psycho-physiological Interaction (PPI) effect and its relationship with participants’ behavioural performance.

PPI is a general linear model with an interaction effect that allows us to investigate differences in FC between two experimental conditions (see Methods). We call such FC that tracks variable cognitive demand effective FC (eFC). Using PPI we found that as cognitive demand increased, eFC increased between the vPCu and IoN and between the dPCu and EoN. However, when the cognitive demand was low the eFC association was reversed (Figure 4b). In other words, the dPCu was more connected to visual and motor networks in difficult (vs. easy) conditions, and more connected to DMN regions in easy (vs. difficult) conditions. On the contrary, the vPCu was more connected to the rest of the DMN and interoceptive regions in difficult (vs. easy) conditions, and more connected with visual and primary sensory networks in easy (vs. difficult) conditions (Figure 5a). As these patterns emerged in both tasks, we averaged the effect across the two tasks to emphasise the common features (brain heat maps on the Figure 5a). An additional hierarchical clustering analysis provided objective evidence that the eFC between the d/vPCu and IoN/EoN were reversed by different levels of difficulty (Figure 5b). The reversed modulation of the v/dPCu’s FC was found after controlling for spurious statistical correlations between their respective time courses (See Methods).

**Figure 5:**
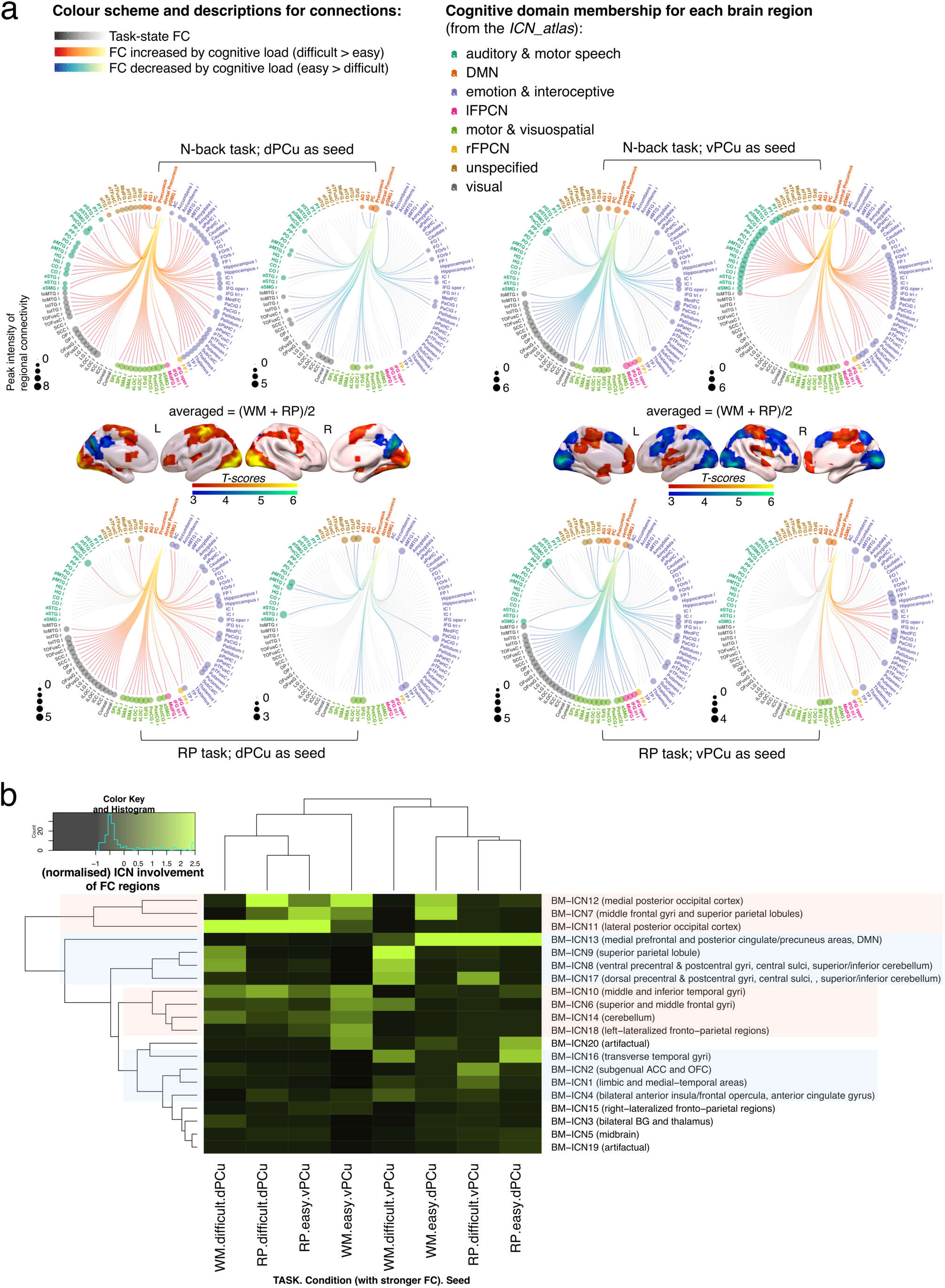
PPI analyses for the two tasks revealed an inverse pattern of task modulated FC of the two seeds (v/dPCu’s), which is common to both tasks. (a) FC overviews for each task are shown in the circular connectogram plots. These were constructed by segmenting T-score PPI maps to anatomical regions as defined by the Harvard-Oxford cortical and subcortical structural atlases in the CONN toolbox (https://web.conn-toolbox.org/). Cerebellar regions are not included in the figures but are reported in the text and tables (see supplementary activation result tables). The *ICN_atlas* toolbox was used to construct the connectograms (full description of this procedure is provided in the supplementary material, sFigure1). The diameter of the circles at the vertices of each connectivity plot indicate the peak intensity (biggest T-score) within this brain region. The whole-brain maps in the middle row depict averaged results between two tasks. Warm colours on the brain maps indicate the regions whose FC with the seed is increased upon an increased cognitive demand (difficult > easy), while cold colours indicate the regions who FC with the seed is decreased upon an increased cognitive demand (easy>difficult). (b) Hierarchical clustering of the v/dPCu’s PPI results confirms the interplay between v/dPCu’s task-modulated FC and internally/externally oriented networks. The brightness values in the heatmap represent normalised ICN spatial involvement (normalised across the 20 network domains per seed per condition per task). The distribution of ICN spatial involvement is shown in the top left corner. To visually identify patterns, the rows and columns of a heatmap are sorted by hierarchical clustering trees. Across columns, the regions having strengthened FC with the v(d)PCu are clustered with the regions having weakened FC with the d(v)PCu upon increased cognitive demand. Across rows, clustered networks form two main categories, which respectively were the internally-oriented networks such as the DMN, subgenual ACC, AIC and OFC, and the externally-oriented networks such as the visual networks, superior and middle frontal gyri and FPCNs.

Further, we found that the positive eFC between vPCu and IoN, and between dPCu and EoN, could predict the increased respond time (RT) that participants needed to deal with increased task difficulty. Specifically, the eFC between the dPCu and FPCN regions (including inferior frontal gyrus, medial frontal gyrus and superior temporal gyrus) was significant in this analysis as was the eFC between the vPCu and DMN regions (including AnG, PCu and precentral gyrus) (Figure 6). It is noteworthy that eFC predicted task performance while the contrast-based activation did not.

**Figure 6:**
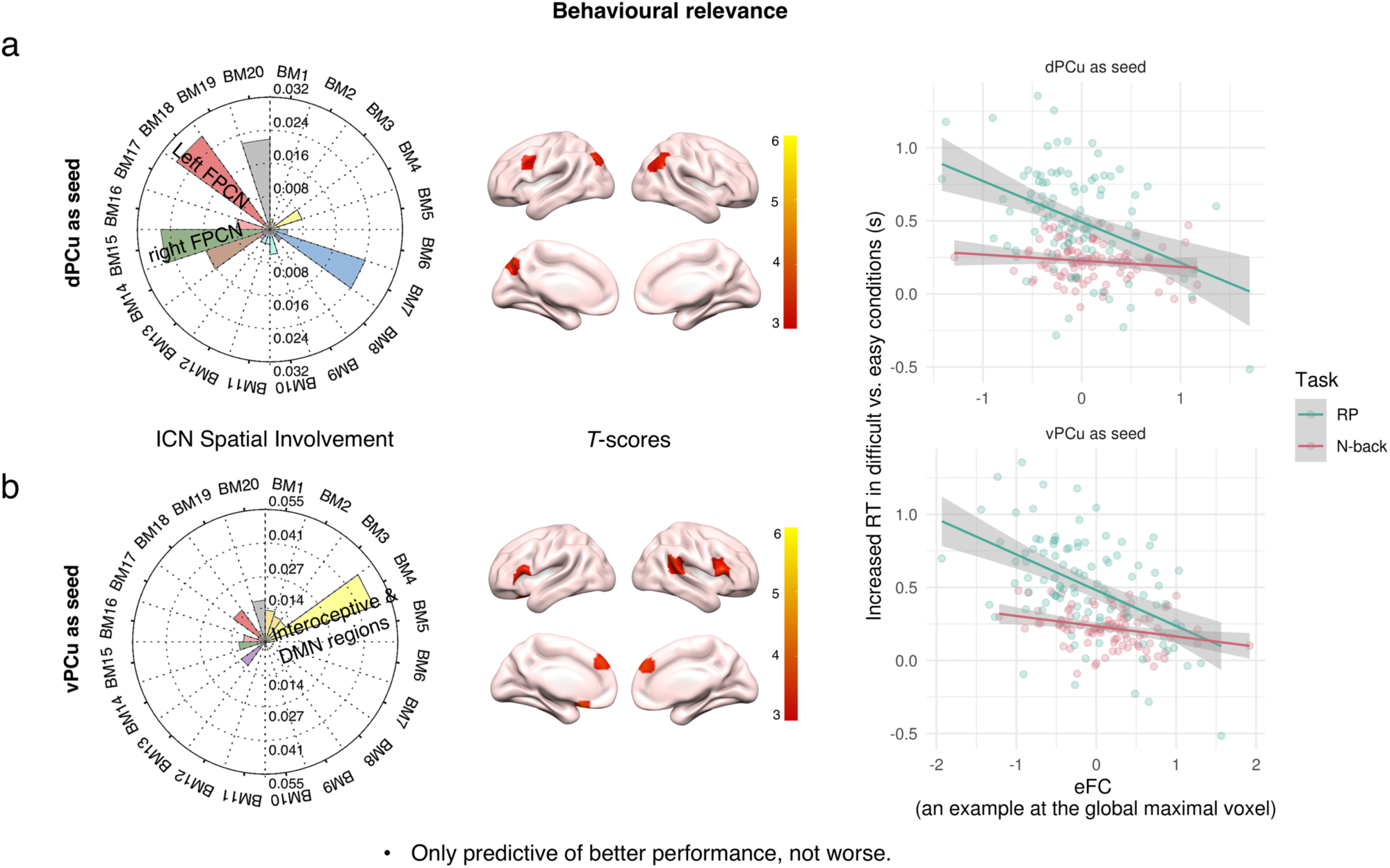
Task modulation of the d/vPCu’s FC (i.e. eFC) from the PPI analysis predicts reaction time of correct responses. The stronger the dPCu’s FC with bilateral frontoparietal regions, the quicker the participant responds correctly in the difficult condition (using their RT in easy condition as a baseline). Plots show the linear relationships between eFC and (relative) reaction time to correct responses at the global statistical maxima of the two analyses. BM-ICN involvement of the significant regions is demonstrated using the *ICN_atlas* toolbox (also see figure 3).

### The causal influence from dPCu→vPCu and from vPCu→dPCu was differentially modulated upon low and high cognitive demand

Our findings so far suggest that the v/dPCu are respectively focused on internal and externally relevant information processing. To obtain further evidence that the PCu is the brain region where these two types of information get integrated, we explored the directionality of information flow between the vPCu and dPCu and its modulation by task demands.

To achieve this, we adopted the dynamic causal modelling (DCM) framework which infers evidence for causal connectivity between regions upon external perturbation during a task (K. E. Stephan et al., 2010). According to our hypothesis, we specified 11 possible model structures, based on how the task difficulty might modulate causal connectivity between the vPCu and dPCu (Table 4).

**Table 4:** Model comparison result suggests that the causal influence from the vPCu to dPCu is modulated in easier conditions and then swapped from the dPCu to vPCu in more difficult conditions. The first 4 rows/aspects of the model structure specified 11 DCM models. The first four DCMs that adopted “one-state”, “deterministic” neural mass model won over the “two-state”, “stochastic” ones, according to the posterior probability at the group level (assuming a fixed factor effect). Among the “one-state”, “deterministic” DCMs, the third model wins over others with consistently higher posterior probability and higher exceedance of model evidence.

| Model<br>Config. | 1 | 2 | 3 | 4 | 5 | 6 | 7 | 8 | 9 | 10 | 11 |
| --- | --- | --- | --- | --- | --- | --- | --- | --- | --- | --- | --- |
| endogenous EC | V↔D | V↔D | V↔D | V↔D | V↔D | V↔D | V↔D | V↔D | V↔D | V↔D | V↔D |
| modulated EC (diff.) | V→D | D→V | V→D | D→V | V↔D | V→D | D→V | V→D | D→V | V→D | D→V |
| modulated EC (easy) | V→D | D→V | D→V | V→D | V↔D | V→D | D→V | D→V | V→D | none | none |
| neural mass model<br>1 = one-state & deterministic<br>2 = two-state & stochastic | 1 | 1 | 1 | 1 | 2 | 2 | 2 | 2 | 2 | 2 | 2 |
| posterior probability in N-back | 0.00 | 0.64 | 0.35 | 0.00 | 0.00 | 0.00 | 0.00 | 0.00 | 0.00 | 0.00 | 0.00 |
| posterior probability in RP | 0.00 | 0.00 | 0.99 | 0.00 | 0.00 | 0.00 | 0.00 | 0.00 | 0.00 | 0.00 | 0.00 |
| (relative) model evidence in N-back | 0 | 15.51 | 14.93 | 8.27 |  |  |  |  |  |  |  |
| (relative) model evidence in RP | 23.48 | 29.07 | 55.09 | 0 |  |  |  |  |  |  |  |

Bayesian model comparisons among the 11 models showed that, for the micro-circuit neural model embedded in the DCM, the bi-linear, one-state and deterministic model class was better for fitting our data than the non-linear, two-state and stochastic one (Marreiros et al., 2008), with an exceedance of log-evidence (an indication of how good a model is by weighing the goodness of data fitting against the model complexity) > 1*10^16^ in the N-back task and > 5*10^16^ in the RP task (Table 5). By comparing the possible ways of how task difficulty modulated the directed information flow we established two models which had higher model evidence than the others: (1) the “exchange” model, which specifies an effective connectivity (EC) modulation of dPCu → vPCu in the easy condition and vPCu → dPCu in the difficult condition; (2) the “forward” model, which specified the EC of dPCu → vPCu to be modulated in both difficult and easy conditions.

In the RP task, the “exchange” model was identified as the single winning model by the BMS (Figure 7). Although in the N-back task the “forward” model had higher posterior probability than the “exchange” model by 0.3, the magnitude of its model evidence did not allow us to draw a safe conclusion of favouring it over the other model (K. E. Stephan et al., 2010).

**Figure 7.**
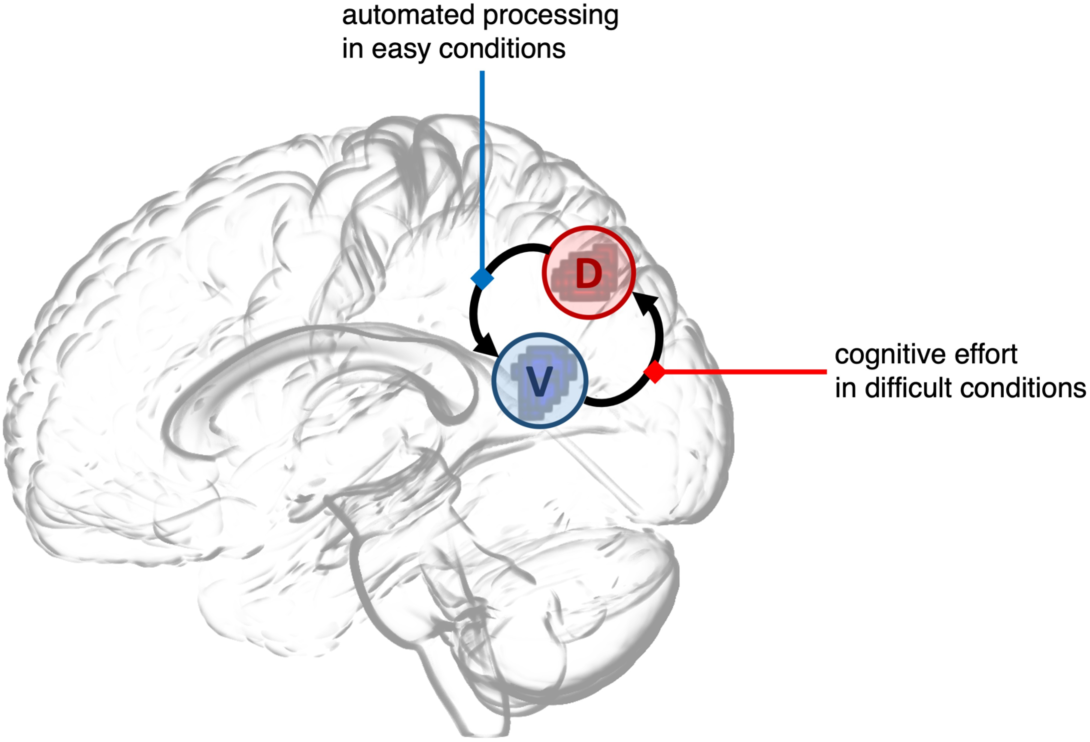
Illustration of the model structure of the winning DCM model.

## Discussion

The present study investigated the functional differentiation of the DMN and proposed a role for the precuneus (PCu) in mediating external and internal information binding. We showed that the PCu has differential connectivity with the rest of the DMN and cognitive control networks. These are not an epiphenomenon of structural or resting-state functional connectivity, but instead they track cognitive demands in a working memory and a high-order relational processing task. In addition, we found evidence for directed interactions from dPCu to vPCu during easier cognitive conditions and from vPCu to dPCu during more difficult conditions. Collectively, these results highlight a causal interaction between the different information processing streams of the d/vPCu for binding internal (e.g., self-related or high-level stored information) and external (e.g., incoming environmental stimuli) information.

Although previous work also hinted at the different roles in the ventral and dorsal parts of the PCu, this is the first systematic demonstration that the v/dPCu are respectively more closely connected to internally and externally oriented networks. A previous resting-state FC study showed that the vPCu is functionally closer to the DMN, while the dPCu is more coupled with the superior occipital lobe and cognitive control regions such as the SPL (Zhang & Li, 2012). A structural connectivity study also showed that from dorsal to ventral parts of the PCu, there is a spectrum of increasing connectivity with DMN regions (such as hippocampus and mPFC) and decreasing connectivity with sensorimotor, visual networks, thalamus and executive regions (such as SPL, prefrontal and premotor areas) (Cunningham et al., 2017).

In a previous task-based study, Leech and colleagues (2011) using a dual N-back experiment found a functional differentiation between the (deactivated) dorsal PCC and the (activated) ventral PCC. The dorsal PCC in their study is close to what we call vPCu in this study. This region (BA 31) lies in the borders between BA 7 (PCu) and BA 23 (PCC) and has been considered to be part of the PCu or PCC by different authors (Cavanna & Trimble, 2006). Leech and colleagues identified distinct FC patterns of ventral PCC and dorsal PCC, the former not only decreasing its positive FC with the DMN but also reducing the negative FC with cognitive control networks (thus being less integrated with the whole brain in general); on the other hand, they found the dorsal PCC increased its connectivity with both the DMN and cognitive control networks, in the difficult versus easy contrast. Noticeably, we did not find any significant negative FC for our PCu seeds. Non-neuronal noise is unlikely the reason for our lack of anticorrelation here, because we have taken rigorous measures to alleviate non-neuronal noise sources, over and above the data-driven denoising utilised in the HCP preprocessing pipeline. Furthermore, there have been studies casting doubts on the nature of the anti-correlation between the DMN and task-positive networks, suggesting that the anti-correlation diminished when global signal regression was not used or high-frequency signals were considered (Caballero-Gaudes & Reynolds, 2017; Craig et al., 2017).

A recent study probing the DMN’s role in auto-pilot behaviour suggested that via the connectivity between PCC/PCu and middle temporal lobe, the brain gains access to stored information and utilises learnt rules to complete the task at hand (Deniz Vatansever et al., 2017). Here by using two levels of difficulty, we also found that when more cognitive effort was needed, stored information was employed. This manifested in the form of increasing connectivity between the vPCu and other parts of the DMN and elevated causal influence from the vPCu to dPCu; meanwhile, the dPCu connected more with attentional systems dealing with increased difficulty and lack of readily available answers.

We propose to interpret our series of findings through the lens of the predictive coding theory (Friston & Kiebel, 2009) (Figure 8). Accordingly, externally driven and internally driven information are exchanged in the PCu and this process is mediated by a “causal loop” between the PCu’s dorsal and ventral subdivisions. By binding internal priors encoded in high-level cortex (gradually shaped from previous experience) and external representations from sensory cortices, the brain acts as Bayesian machine to interpret ambiguous and quickly shifting environmental cues.

**Figure 8:**
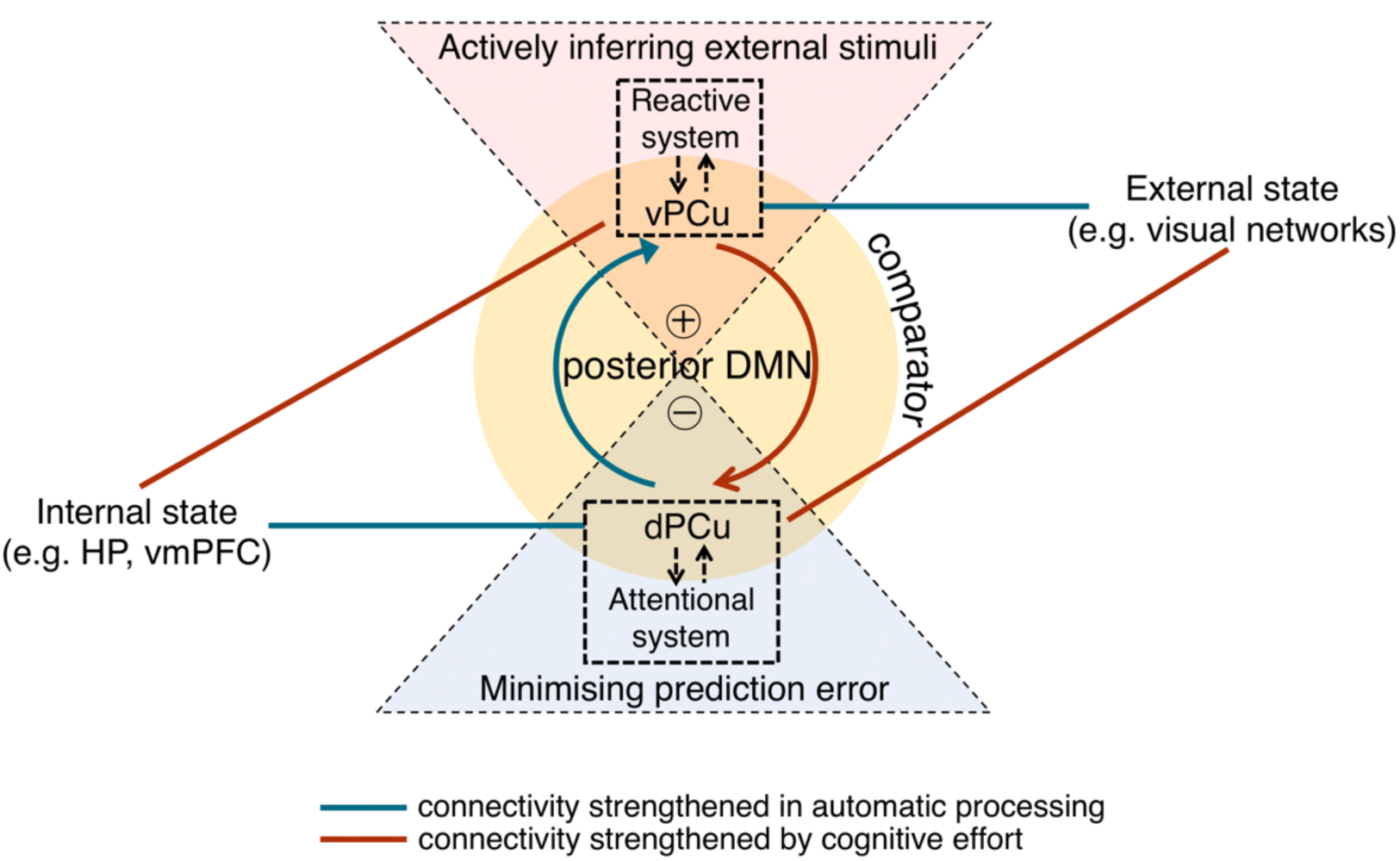
A conceptual predictive model centring at the medial posterior DMN. The medial posterior DMN behaves like a predictive system, with its dorsal part associated with an attentional system (comprising FPCN regions such as the IPS, middle and lateral frontal cortices) and its ventral part associate with a reactive system (comprising regions such as anterior insula) (Tops et al., 2014). The connectivity from each module is derived from our studies, with the pink colour indicating being stronger in difficult conditions, and blue colour indicating being stronger in easy conditions. The (directed) connectivity pattern from our result suggests that external and internal information is exchanged by the reciprocal loop between the vPCu and dPCu. This exchange of information might support the predictive coding theory which emphasises that external information is always inferred (rather than simply being represented from a bottom-up direction) based on an internal belief of the world (prediction).

The functional differentiation within the PCu may just be an iceberg tip in terms of DMN characteristics. Recently, more studies have evaluated the idiosyncratic character of the DMN. Braga and Buckner (2017) proposed the DMN encompasses several networks, the spatial pattern of which differs between individuals. Other studies also suggested that the DMN is a “transmodal” region (Braga et al., 2013), meaning it has many specialised subdivisions. For example, the AG is for attentional control and semantic comprehension (Lyu et al., 2019; Ralph et al., 2017; Seghier, 2013), ACC for value-based perception and error detection (Chudasama et al., 2013; Fleck et al., 2006; Monosov, 2017) and hippocampus for memory encoding (Bird & Burgess, 2008) etc. However, the PCu’s function seems to have an all-encompassing nature since it is activated under all kinds of cognitive demand where other parts of DMN are employed (Laird et al., 2009). Indeed, graph theoretic analyses on the human brain connectome have suggested that the PCu is the central hub of brain connectivity (P van den Heuvel & Sporns, 2011; Tomasi & Volkow, 2011; van den Heuvel & Sporns, 2013). The PCu, besides the DMN, has also been found to be most closely connected to the thalamus (Cunningham et al., 2017). At this brain connectivity hub, information from all sources converges, endowing a narrative construction of reality: It have also been shown that during movie watching the temporal pattern of the PCu’s activity, compared to other brain regions, can track event boundaries of changing scenes in the most abstract level (Baldassano et al., 2017). Based on previous work and our current study, we believe the PCu is central for linking the function of the DMN subdivisions, thus playing a key role in integrating the brain’s information flow from all sources.

Although the PCu was our focus in this study, it is not the only DMN subregion that demonstrates a functional fragmentation. We also observed a similar functional differentiation in the AnG’s activation pattern, but only in the N-back task. Evidence also comes from previous studies which have discussed the activation patterns of the DMN subregions, such as the mPFC (Bzdok et al., 2013; Kuzmanovic et al., 2018), the IPL (Igelström & Graziano, 2017), the PCC (Leech et al., 2011) and the PCu (Cavanna & Trimble, 2006). Taken together, the evidence suggests that these regions may generally have a ventral (anterior)-dorsal (posterior) gradient of functional division. We therefore postulate that the DMN might comprise ventral/anterior and dorsal/posterior subdivisions that act as interfaces between self (internal)- and environmental-related processing. Our study is agnostic about possible functional subdivisions of other DMN regions besides the PCu, but future studies should continue the exploration of the functional differentiation between and within the DMN subregions. The DMN has been postulated to be at the highest level of information processing in the brain (Margulies et al., 2016) and has the highest number of optimal inter-network connections in the brain (Pappas et al., 2020), along with our findings presented here, we propose that this topological division of internal and external related functions may be a necessary consequence of developmental optimisation of information processing. By continuing this line of investigation, we might gain important insights into the brain’s functional organisation.

## Materials and Methods

### NeuroSynth Meta-analysis of fMRI studies

When this work was carried out, the latest Neurosynth database contained 14,371 neuroimaging studies (https://github.com/neurosynth/neurosynth-data), associated with more than 3,200 text-based features and over 410,000 activation peaks that span a wide range of published neuroimaging studies. Since our focus is DMN functionality during tasks we searched for activation coordinates associated with attentional and executive tasks. These were generated by interrogating the database with the text-based search: “attention* or execut*”. Both forward and reverse inferences were performed in order to assess both necessity and sufficiency of a region’s response to a certain type of task. Specifically, the forward-inference search relies on the probability of observing activation given the presence of the term [i.e., P(Activation|Term)], thus essentially testing the consistency of a region’s activation to a type of task. On the other hand, reverse inference relies on the probability of the specified term being frequently discussed alongside a specific activation [i.e., P(Term|Activation)], thus showing the regions that are selectively associated with the term. Key parameters were based on the default values in the publicly available *NeuroSynth* toolbox (https://github.com/neurosynth/neurosynth). For example, a frequency cut-off of 0.001 for article words was used to determine if a study used the term incidentally or purposely. Resulting activation was FDR-corrected for multiple comparisons using a whole-brain FDR threshold of 0.01 at the voxel level. This generated Z-score maps with values generally bigger than 3. For a better visualisation of the result, we used a cut-off of Z=4 to only show bigger clusters.

### FMRI & Diffusion-Weighted Data Analyses

#### Tasks selected

We selected the Relational Processing (RP) task and the N-back working memory (N-back) task from the human connectome project (HCP), as these have two levels of difficulty, therefore affording the possibility to infer the brain’s response to varying cognitive demand. Analyses for the two tasks were conducted independently first, and then results were averaged across the two tasks to emphasise domain-general brain activity. In the main text, domain-specific terms indicating a certain level of specific cognitive demand (such as 0-back and 2-back in the N-back task) were substituted with the general terms of “easy” and “difficult” to indicate the load of general cognitive abilities.

#### Image preprocessing

The ready-to-use HCP data has already been minimally preprocessed and quality-checked by the distributors (Glasser et al., 2013) and we carried out extra preprocessing steps with SPM12 (http://www.fil.ion.ucl.ac.uk/spm). Specifically, the data was smoothed with a Gaussian kernel of 6 mm FWHM (full-width half maximum) and no low-pass filtering was used as it might reduce signal strength and sensitivity. No global signal regression was used, for it may cause anti-correlation artefacts by shifting the distribution of FC towards negative values (Chai et al., 2012; Murphy et al., 2009). To reduce the influence of non-neuronal confounds, eigenvalues were extracted from BOLD (Blood Oxygen Level Dependent) signals within cerebrospinal fluid (CSF) and white matter template masks and were regressed out from target signals during statistical testing.

#### Activation studies

Whole-brain activation was estimated using the standard SPM GLM approach. Only the trials with correct responses were explicitly modelled. Individual-level GLMs were modelled with difficulty level (2 levels) as the main effect, i.e. difficult (2-back or relational) & easy (0-back or match) conditions, with neuronal nuisance (i.e. CSF, white matter signals), session effects and 6 movement regressors as covariates. Group-level GLMs were then constructed to test the significance of the activation in the population level, with age and sex as covariates.

#### Selection of Regions of Interest (ROI)

We were interested in the functionality of the PCu during attentional demanding goal-directed tasks; therefore, ROIs were selected based on our activation results from the N-back and RP tasks. Both activated and deactivated PCu regions associated with increased level of difficulty of the tasks were considered. To ensure the PCu region that we are considering is actually part of the DMN, we also superimposed it on the DMN canonical mask as defined by the *Conn* network atlas. The seed regions were selected by computing the spatial intersection of our task-related (de)activation clusters (exceeding the cluster-level family-wise-error/FWE-corrected P of 0.05), the PCu as defined by the Conn atlas and the DMN spatial localisation identified by the *Conn* atlas (Whitfield-Gabrieli & Nieto-Castanon, 2012). These selection criteria generated dPCu and vPCu seeds respectively for the N-back and the RP task. Timeseries were extracted from seed regions, and their FC during the course of the experiment was calculated for the two tasks separately. The seeds we used for structural connectivity and resting-state FC analyses were constructed by computing the overlap between the PCu seeds from the two tasks (Figure 2a).

#### Functional connectivity studies

##### Resting-state Seed-based FC of vPCu & dPCu

Using SPM12, we calculated the FC as the partial correlation of the timeseries extracted from the seed regions (v/dPCu) with that from the rest of the brain, after controlling for the effect of non-neuronal confounds (estimated from white matter and CSF) and head movements. Baseline FC and differences between the FC of the (v)dPCu at the group level were estimated with one-sample and paired T-tests.

##### Structural Connectivity of vPCu & dPCu

The acquisition, preprocessing the diffusion MRI (dMRI) images from the HCP and the generation of diffusion tensor maps has been detailed in published articles (Sotiropoulos et al., 2013). Based on the diffusion tensor images we built probabilistic tractography in FSL5 (https://fsl.fmrib.ox.ac.uk/fsl/fslwiki/FDT). The structural connectivity matrix for each individual was obtained using vPCu or dPCu as the seed and the grey matter as the termination mask. Each entry in the connectivity matrix stands for the number of streamlines (out of 5000) from each voxel of the seed map (vPCu or dPCu) that had a 50% change or greater of reaching the grey matter (curvature threshold = 0.2), which was also corrected for the distance/length of the pathways.

A seed’s structural connectivity to the grey matter was calculated by averaging the streamline connections from all the voxels of the seed. This matrix was then projected to a standard MNI brain space for further statistical parametric mapping analyses in SPM12. Similar to FC analyses, baseline and difference of grey-matter connections of the dPCu and vPCu at the group level, were calculated with one-sample and paired T-tests. To brain maps of the statistical parametric values for each ROI were smoothed with a Gaussian kernel of 6 mm FWHM to increase normality of the image data.

##### Task-state Seed-based FC of vPCu & dPCu

Task-state functional connectivity (tsFC) was derived based on partial correlations of the timeseries between the seed and the rest of the brain, after controlling for the effects of the event-related BOLD response (boxcar experiment design convolved with the canonical HRF) weighted by contrasts, as well as other non-neuronal confounds during the course of the experiment.

##### Psycho-physiological interactions (PPI)

PPI was used to evaluate by what amount the cognitive variables during tasks upregulate or downregulate the FC between the seed region and its functionally-coupled regions (Figure 4b).

The implementation of the PPI was similar to the above seed-based FC during tasks, but additionally the GLM model included an interaction term, i.e. a new variable created by dot multiplying the task-related BOLD response and the seed’s timeseries. Individual-level PPIs were estimated separately for the N-back and the RP task, but the final results were averaged across tasks because we wanted to emphasize the common emerging patterns. Parameters of the modulation effect measured by individual-level PPIs were modelled as the dependent variable in a group-level GLM, with the individuals’ RT difference between two conditions as the main regressor, and with their age, sex and task identity as covariates of no interest. RT was included in this analysis to investigate the behavioural relevance of the PPI modulation.

We first conducted PPI analyses independently for the two tasks and we observed a common pattern of the modulation effect between the N-back and the RP tasks. To emphasise that common pattern, we then averaged the individual-level modulatory effects between the two tasks, and conducted another group inference based on the individual averaged beta values. Results of task-specific PPIs and task-averaged PPI are both provided.

##### Anatomical labels and ICN identification based on significant clusters

To make inferences about the cognitive function of significant regions, we used the functionally defined ICN-BM network atlas (https://www.nitrc.org/projects/icn_atlas/) for identifying the intrinsic (functional) connectivity networks (ICN)s involved in the two tasks. The advantage of using the ICN-BM atlas was the nomenclature used not only corresponds to the well-known canonical resting-state connectivity networks, but also to task-based co-activation networks which were generated from a meta-analyses using the BrainMap (BM) dataset (Cole et al., 2016; Smith et al., 2009). This robust correspondence between ICNs and cognitive functions allowed us to make inverse inferences, that is to infer implicit cognitive processes from engaged brain areas.

We used the *Matlab*-based “*ICN_Atlas*” toolbox to determine each significant cluster’s region/ICN correspondence by rating their spatial overlaps with the pre-defined regions/ICNs (Cole et al., 2014, 2016; Ito et al., 2017; Kozák et al., 2017; Laird et al., 2011, 2013; Smith et al., 2009). When reporting the PPI result, we also applied the same strategy to establish the correspondence of the *Conn* atlas and the ICN-BM atlas for visualising both the seed-based FC regions (defined by the *Conn* atlas) and the ICNs they belong to. The correspondence of the two atlases is presented in the Supplementary Materials.

The spatial overlap reported in the main article was calculated as the ratio of the number of activated voxels over the region/ICN volume which the voxels belong to (Kozák et al., 2017). Other ways of representing the spatial overlap, such as calculating spatial overlaps by taking into account the effect size of each significant voxel, were found to generate similar results and are presented in the Supplementary Materials.

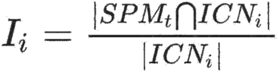

ICN_i_ Spatial Involvement (I_i_): ratio of the number of activated ICN_i_ voxels to ICN_i_ volume

Besides the activation maps and their associated ICN domains shown in the main text, we also report activation tables with detailed coordinates in the Supplementary Materials. For identifying the anatomical regions based on the coordinates, we adopted the toolboxes of *GingerALE* (http://brainmap.org/ale/) and *Talairach Daemon* (http://www.talairach.org/daemon.html) where the Brodmann areas were identified from.

#### Group analysis and multiple-comparison correction

For all statistical parametric mapping analyses, random factor effects (RFX) were used (random effect being the intercept of within-subject GLM fitting) and inferences were made at the group level to allow generalisation to the population. For the group level inference, individual weighted beta coefficient maps were fed into a one-sample t-test that tested for the significance of group means of the factor of interest. The multiple comparison problem was dealt using cluster-extent thresholding within the Gaussian random field framework (*Human Brain Function*, 2004) implemented in SPM. Clusters were usually defined by a default primary voxel threshold of 0.001 (uncorrected). However, more stringent voxel-level thresholds were sometimes used because the statistical power in this study was very high and the conventional cluster-forming threshold of P-uncorrected = 0.001 resulted, on some occasions, on clusters that were too extensive to be anatomically meaningful. In such cases, it has been suggested in the previous literature to use more stringent voxel-level threshold for making sensible inferences (Woo et al., 2014). Detailed threshold values used are reported in the results tables in the Supplementary Materials. The family-wise error rate was controlled at the cluster level, and a threshold of P-corrected < .05 was used to determine significance among clusters.

#### Dynamic Causal Modelling (DCM) specification

Dynamic causal modelling (DCM) is a generative model in a Bayesian framework for inferring hidden neuronal states from observed fMRI measurements (K. E. Stephan et al., 2010). The causal influence estimated from the DCM is neurobiologically interpretable and describes the effect on the strength, or the rate of change, of synaptic connections among neuronal populations, as well as their context-sensitive modulation upon external perturbation during tasks (K. E. Stephan et al., 2010). Using SPM 12, our DCM model specified the endogenous connectivity between vPCu and dPCu to be bi-directional; and on top of that we modelled all possible configurations of how the task difficulty might influence the endogenous connectivity (Table 1). Since DCM is a generative model, it has to make assumptions about local neural populations in each region, and there are different kinds of neural mass models to choose from. In one-state models, each region is assumed to be composed of only single-state neurons, whose activity decays following a perturbation by exogenous input or incoming connections. In two-state models, each region is composed of both excitatory and inhibitory neural populations, the interaction of which can generate more complex dynamics (Marreiros et al., 2008). And stochastic models, compared to deterministic models, take into account neural noise for the local neural mass model (Daunizeau et al., 2012).

Our focus was on the directionality of connectivity at the regional level; therefore, we compared model structures representing all four combinations of how the two conditions (difficult vs. easy) effected the mutual directional connectivity between d/vPCu. Based on that, we tried both the one-state, deterministic (the default) and the two-state, stochastic DCM class for modelling local neural dynamics.

#### DCM estimation

To determine the most likely model structure, we applied a fixed factor effect (FFX) Bayesian model selection (BMS) procedure to all 11 models estimated across all participants independently for the N-back and the RP task. The FFX was used as opposed to a random factor effect (RFX) because we hypothesised the mechanism to be general across all subjects. Finally, the model with the highest model evidence was selected over others. Model evidence is the probability of obtaining the observed data given a particular model (Klaas Enno Stephan et al., 2009).

Upon the selection of the best model structure, individuals’ effective connectivity (EC) parameters were then used to predict their behavioural performance (reported in the Supplementary Materials). To do so, linear regressions were conducted to probe the relationship between the RT and modulation of EC independently for the EC from dPCu to vPCu and the EC from vPCu to dPCu, with confounding covariates of age, sex and the mean activity of the vPCu and dPCu during tasks. The modulatory effect of task demand was estimated as the DCM parameters in the B matrix (https://www.fil.ion.ucl.ac.uk/spm/doc/spm12_manual.pdf). These parameters and RT were logarithmic transformed to be more normally distributed.

